# Integrative Analysis the characterization of peroxiredoxins in pan-cancer

**DOI:** 10.1101/2020.10.09.334086

**Authors:** Lei Gao, Jialin Meng, Chuang Yue, Xingyu Wu, Quanxin Su, Hao Wu, Ze Zhang, Qinzhou Yu, Shenglin Gao, Song Fan, Li Zuo

## Abstract

Peroxiredoxins (PRDXs) are antioxidant enzymes protein family members that involves the process of several biological functions, such as differentiation, cell growth. Considerable evidence demonstrates that PRDXs play critical roles in the occurrence and development of carcinomas. However, a systematic analysis of PRDXs in cancers is deficiency. Therefore, we perform a comprehensive analysis of PRDXs in 33 cancer types including mRNA expression profiles, genetic alterations, methylation, prognostic values, potential biological pathways and target drugs. Moreover, we validated that PRDX6 could regulate cancer cell proliferation via JAK2-STAT3 pathway and involve into the process of cell cycle in bladder cancer.

## Introduction

Peroxiredoxins (PRDXs) are a large group of antioxidant enzymes protein family with more than 3500 members^1^, owning a cysteine residue that involve into the process of reduction of peroxides^2^, which are widely expressed in almost all organisms^3^. Based on the number and location of the active cysteine residues and other factors, PRDXs can be classified into three subfamilies: typical 2-Cys, atypical 2-Cys, and 1-Cys^4, 5^. According to the structural information around active sites, PRDXs are also divided into six subfamilies: AhpC-Prx1, BCP-PrxQ, Tpx, Prx5, Prx6, and AhpE^6^. At present, there are 6 known mammalian PRDX members, named PRDX1, PRDX2, PRDX3, PRDX4, PRDX5, PRDX6^7^. The family members are involved in several biological processes, such as cell differentiation, metabolism^8^, inflammation^9^, cellular protection against reactive oxygen species (ROS) ^10^, embryonic development and cellular homeostasis^11^.

Although the cardio- and neuroprotective effects have been well established^12, 13^, increasing evidences demonstrate that PRDXs play critical roles in the process of carcinogenesis and the development of drug resistance. The function of PRDX1 was first reported to act as an antioxidant enzyme because of its high susceptibility to oxidative stress^14^, which plays important roles in cell growth, differentiation and apoptosis^15^. PRDX2 is association with the proliferative, migratory and metastatic activities of melanoma^16^. PRDX3 has been demonstrated a higher expression in tumor tissues of breast cancer, cervical cancer, liver cancer and prostate cancer than the normal tissues, and is related to the aggressive phenotype of these tumors^17-20^. PRDX4 could increase the proliferation rate of prostate cancer cell and promote the metastasis of ovarian cancer, breast cancer, lung cancer and oral squamous cell carcinoma^21-24^. It was also reported that highly expressed PRDX5 was associated with short overall survival (OS) in patients of ovarian cancer^25^. In breast cancer, the level of PRDX6 is elevated, and associated with an invasive phenotype, PRDX6 caused metastatic potential was also observed in prostate cancer^26, 27^. After searching of literatures, we found that the expression levels of PRDX1, PRDX3 and PRDX6 were up-regulated in cisplatin resistance status for erythroleukemia, breast cancer and ovarian carcinoma^28, 29^. However, the comprehensive function of PRDXs in pan-cancer is unclear till to now.

In order to systematically investigate the function of RPDXs in diverse tumor types, the mRNA expression profiles and their relationship with OS situation among 33 cancer types based on the TCGA database were explored, as well as the association with tumor microenvironment, genetic alteration and drug response activity. Considering that ROS are implicated as critical mediators of tissue damage in patients with bladder cancer (BLCA)^30^ and systematically analyses of PRDXs in BLCA were rare. Then comprehensive analyses of PRDXs and BLCA were performed. Moreover, the effects of aberrantly expression levels of PRDX6 in cell growth, apoptosis and the potential mechanisms were verified in T24 and TCCSUP cells.

## Results

### The gene expression and (overall survival, OS) of PRDXs in pan-cancer

To investigate the expression pattern and prognostic value of PRDXs across diverse cancer types, we collected the clinical information of 33 tumors and corresponding cancer tissue RNA-seq data from the TCGA database. An obvious difference regarding the expression levels of the six PRDX genes were observed (**Figure 1A**). The expression pattern of PRDXs in different tumors were variable, PRDX1, PRDX2, PRDX4 and PRDX5 were mainly up-regulated expression in more than 10 cancer types. However, PRDX3 and PRDX6 were primarily down-regulated expressions in cancers (**Figure 1B, Supplementary Figure1**). Next, according to OS of cancer patients, univariate Cox proportional hazard regression methods was utilized to explore the relationship between PRDXs and OS. The results demonstrated that PRDX4 and PRDX6 were associated with the OS of 8 cancer types (ESCA, KICH, KIRP, LAML, LGG, LIHC, MESO, UVM) and 9 cancer types (ACC, BLCA, HNSC, KIRP, LGG, LIHC, OV, THYM, UCEC), respectively. While the rest PRDXs were only related with the OS of less than 7 cancer types. In additional, we found that PRDX2, PRDX3 were commonly associated with poor survival, PRDX1, PRDX4 with favorable survival, while the direction of the correlation varied depending on the cancer type for PRDX5, PRDX6 (**Figure 1C, Supplementary Table1**). In more detail, PRDX5 predicted poor prognosis for LAML, UVM, but predicted advantage survival for KIRP, LGG, OV. PRDX6 was related with poor prognosis of patients with ACC, BLCA, HNSC, KIRP, LGG, THYM and UCEC, but indicated favored survival for patients with OV.

**Figure 1.**
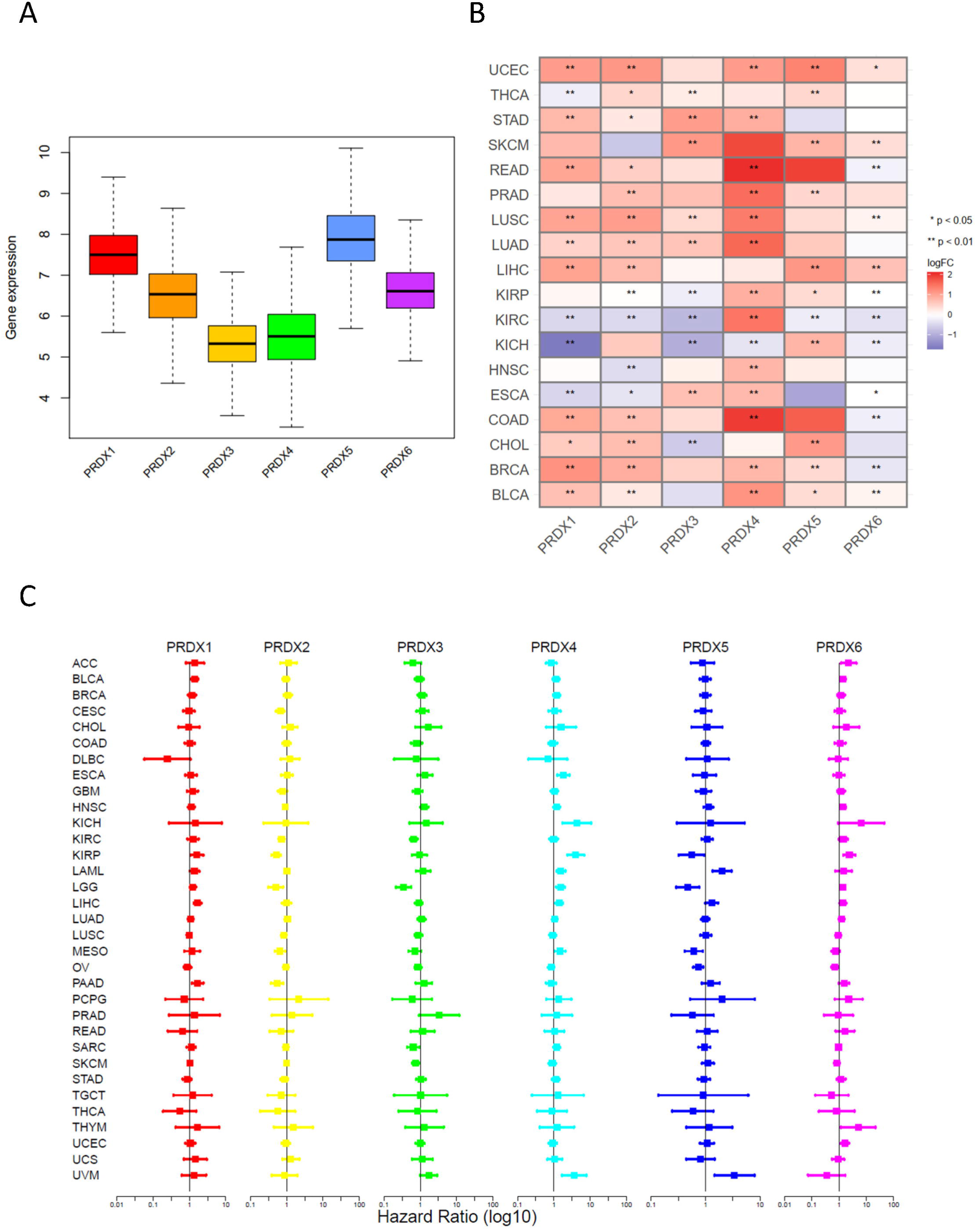
Expression levels of PRDX genes and association with overall survival in pan-cancer. (A) Boxplot to display the distribution of PRDX genes expression across all 33 type tumors. (B) Heatmap to show the difference of PRDX gene expression comparing tumor to normal samples based on log2(fold change) for 18 tumor types. (C) The forest plots for overall survival with hazard ratios (log10) and 95% confidence intervals for 33 different cancer types.

### Genetic alterations and methylations of PRDXs in pan-cancer

Genetic alterations of PRDXs were identified in the different cancers from cBioprotal website. As for the cancer type level, the top-five tumors with PRDXs genetic alterations were Endometrial Carcinoma, Ovarian Epithelial Tumor, Bladder Urothelial Carcinoma, Cholangiocarcinoma, Esophageal Squamous Cell Carcinoma (**Figure 2A**) and the mutation distribution of each PRDX was displayed in **Supplementary Figure2**. Then the copy number variation (CNV) situations and methylations of PRDXs in pan-cancer were explored based on GSCA Lite. The results demonstrated that the mRNA expressions of PRDXs was mainly positively correlated with CNV (**Figure 2B**), negatively with methylation (**Figure 2C-D**), with only a few had no significant, which suggested that PRDXs might be the CNV and methylation drive genes.

**Figure 2.**
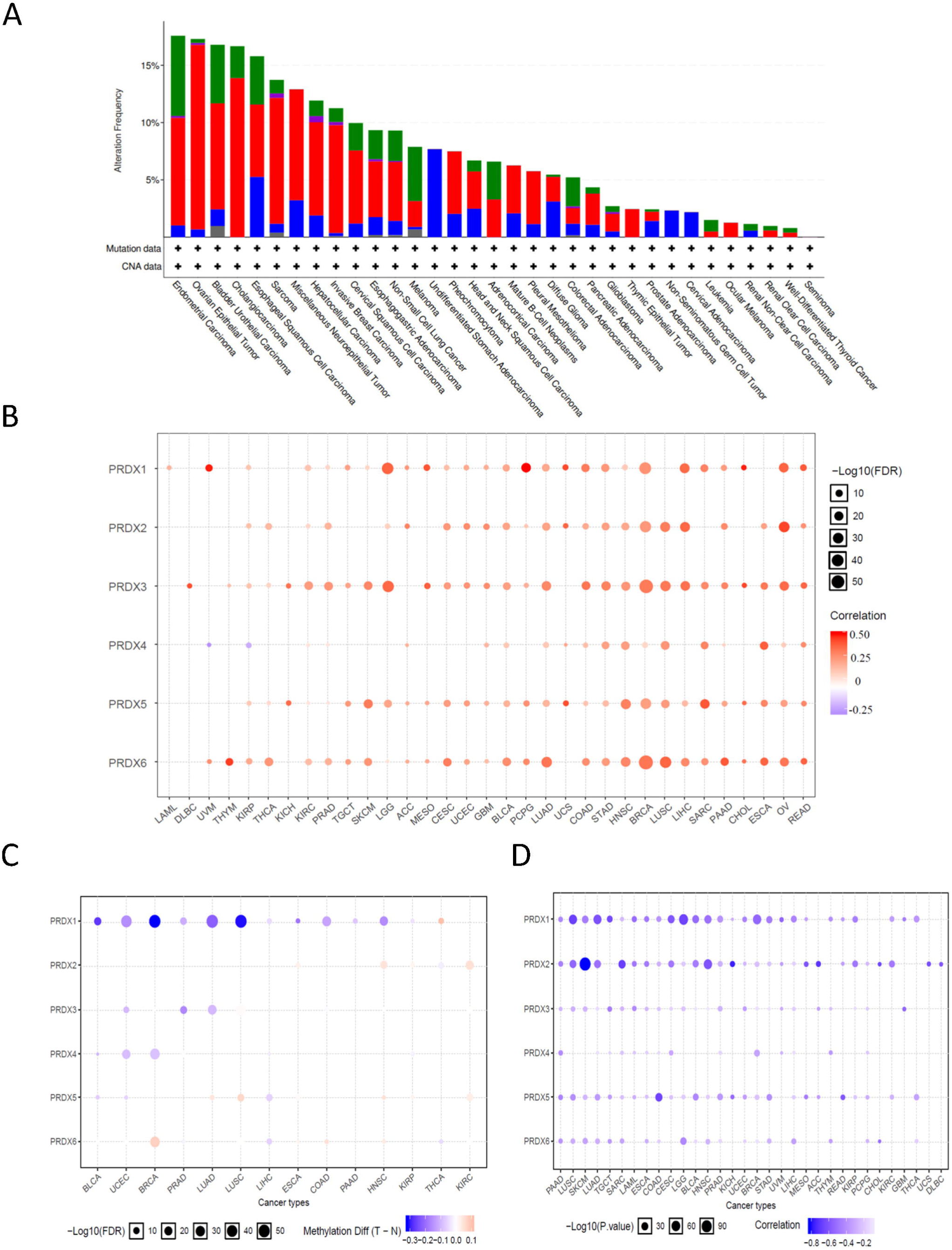
The genetic alterations of PRDXs and associations with mRNA expression. (A) Total genetic alterations of PRDX genes based on cBioportal website. (B) Bubble map indicated the relationship between PRDX mRNA expression and CNV. Red dots indicate that gene CNV levels were positively associated with mRNA expression, blue dots indicate the negative association. (C) The differential methylation of PRDXs between TCGA cancer and normal tissues. Blue dots show methylation down-regulation in tumors, and red dots show methylation up-regulation in tumors. (D) Associations between DNA methylation and PRDXs expression.

### Associations between PRDXs and immune subtypes, tumor stemness, drug sensitivity and signaling pathways

Six subtypes of immune infiltrates, C1 (wound healing), C2 (INF-r dominant), C3 (inflammatory), C4 (lymphocyte depleted), C5 (immunologically quiet), and C6 (TGFβ dominant) were obtained from the study of David et al^31^. Correlations between higher levels of PRDX1, PRDX4, PRDX6 and type 1, 2, and 6 infiltrates (C1, C2 and C6), suggesting the tumor promote roles of these PRDXs (**Figure 3A**). On the contrary, PRDX2 and PRDX3 had higher expression in C4 subtypes, indicating the suppressive roles of them. Considering that PRDXs were associated with drug resistance and drug resistance were closely related with tumor stemness^32^, the relationship between PRDXs expression profiles and tumor stemness were analyzed. The results showed that PRDXs were strongly associated with the tumor stemness across cancer types (**Figure 3B, Supplementary Table 2**). Moreover, the correlation between tumor stemness and TGCT high to 0.66 (*P* < 0.001). Then the relationship between PRDXs and 10 cancer-related pathways were analyzed based on GSCALite website (**Figure 3C**). The results showed that PRDX1 had significantly activation effect of apoptosis, cell cycle, while inhibitory of EMT and RTK. PRDX2 had significantly activation effect of apoptosis, cell cycle, DNA damage response and Hormone AR, while inhibitory of EMT and RTK.

**Figure 3.**
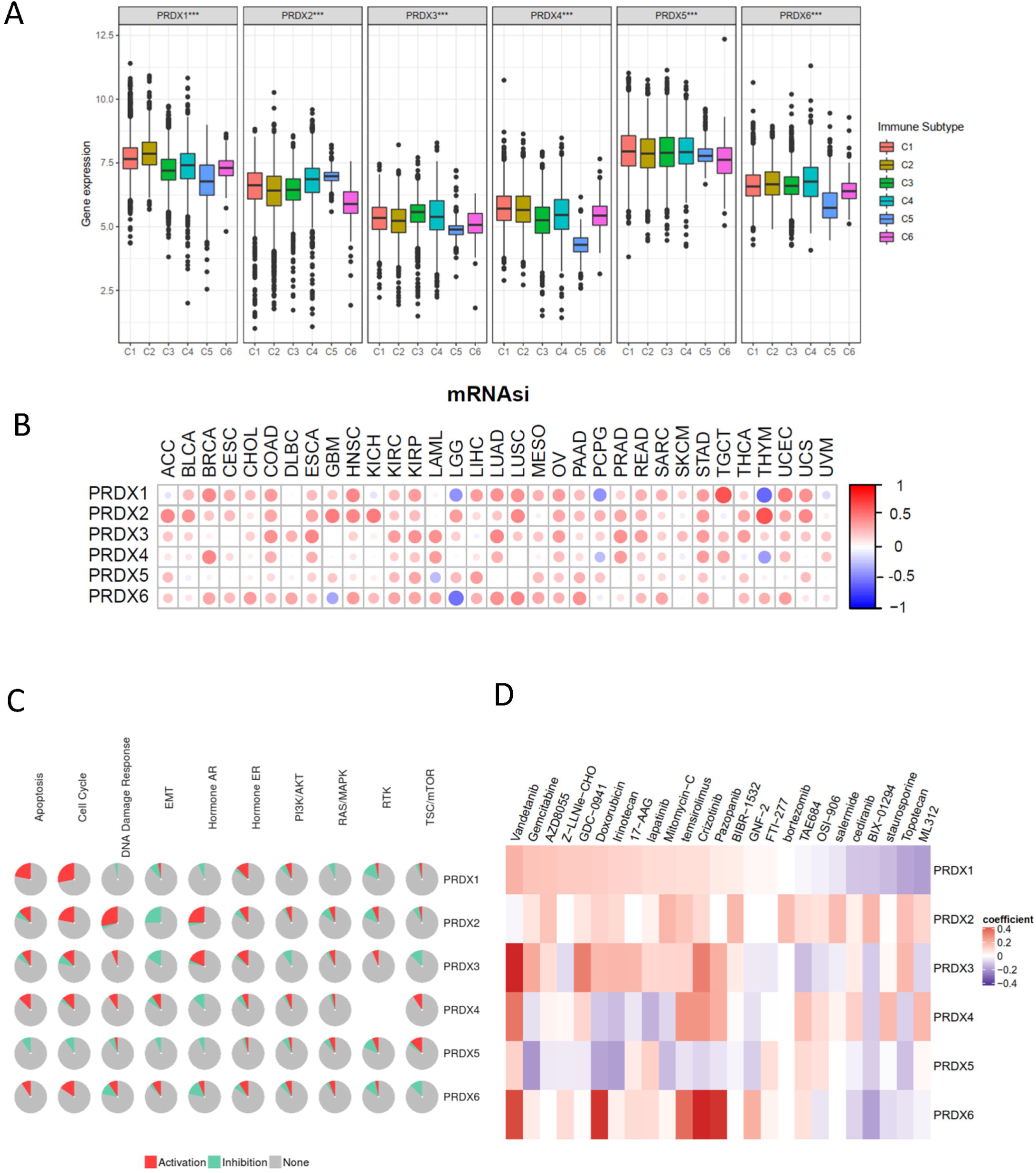
The immune infiltrate subtypes, cancer stemness, pathways and sensitivity to drugs of PRDXs. (A) Association of PRDX gene expression with immune infiltrate subtypes among all the tumor types tested by ANOVA. (B) Correlation plots of the relationship between PRDX gene expression and tumor stemness of 33 diverse tumor types. (C) Fan diagrams show the percentage distribution of PRDXs’ function (activation or inhibition) for related pathway in all cancers. (D) Heatmap of drug sensitivity / tolerance of PRDXs with high expression.

Overall, PRDX1, PRDX2, PRDX3, PRDX4, and PRDX6 had significantly activation effect of apoptosis, cell cycle, while PRDX5 had strongly inhibitory of apoptosis and cell cycle. Next, the five most potential cancer drugs for each PRDX members (p<0.05, standardized coefficient > 0.1) were analyzed based on PharmacoDB^33^ database (**Figure 3C, Supplementary Table 3**). The results demonstrated that pazopanib, vandetanib, lapatinib and cediranib were associated with PRDX1, PRDX3, PRDX4, PRDX6, which suggested that the PRDXs may be strongly associated with the EGF and VEGF signaling pathway. Moreover, Gemcitabine, Doxorubicin, AZD8055 (mTOR inhibitor), GDC-0941 (PI3K inhibitor), Irinotecan (cytotoxic chemotherapy drug), 17-AAG (HSP90 inhibitor), Mitomycin-C (cytotoxic), and temsirolimus drugs were strongly associated with PRDXs (**Figure 3D**).

### Expression profiles, genetic alterations of PRDXs and the correlations with immune subtypes, stages in bladder cancer

Considering that the relationship of ROS and tissue damage in BLCA patients, and the roles of PRDXs in ROS, a thorough evaluation of PRDXs function in BLCA is quite necessary. The mRNA expression, genetic alterations were analyzed. The PRDXs all highly expressed in BLCA tumor tissues as compared with normal tissues (**Figure 4A**), though there is no significant different of PRDX2, PRDX3 between tumor and normal tissues. The genetic alterations of PRDX1-6 were 4%, 1.6%, 1.6%, 3%, 2.4%, 13%, respectively (**Figure 4B, D**), mainly were CNV mutation (**Figure 4E**). The methylation levels of PRDX2-6 between tumor and normal tissues were significantly decreased (*P* < 0.001) (**Figure 4C**). Moreover, the mRNA expression of PRDXs were strongly associated with CNV and methylation in tumor tissues (**Figure 4E**), which were consistence with the analyses of pan-cancer (**Figure 2B-D**). Then the analyses of immune subtypes showed that PRDX1, PRDX4 and PRDX6 were strongly related with type 1, 2 infiltrates (C1, C2), and lower expression in C3 subtypes, indicating the potential tumor promoters, which were similar with the results of pan-cancer (**Figure 5A**). In additional, the mRNA expressions of PRDX1, PRDX4 and PRDX6 were closely related with tumor stage of BLCA (**Figure 5B**). The immunohistochemistry (IHC) results obtained from the Human Protein Atlas (HPA) database showed that PRDX1-6 were high expressions (**Figure 5C**).

**Figure 4.**
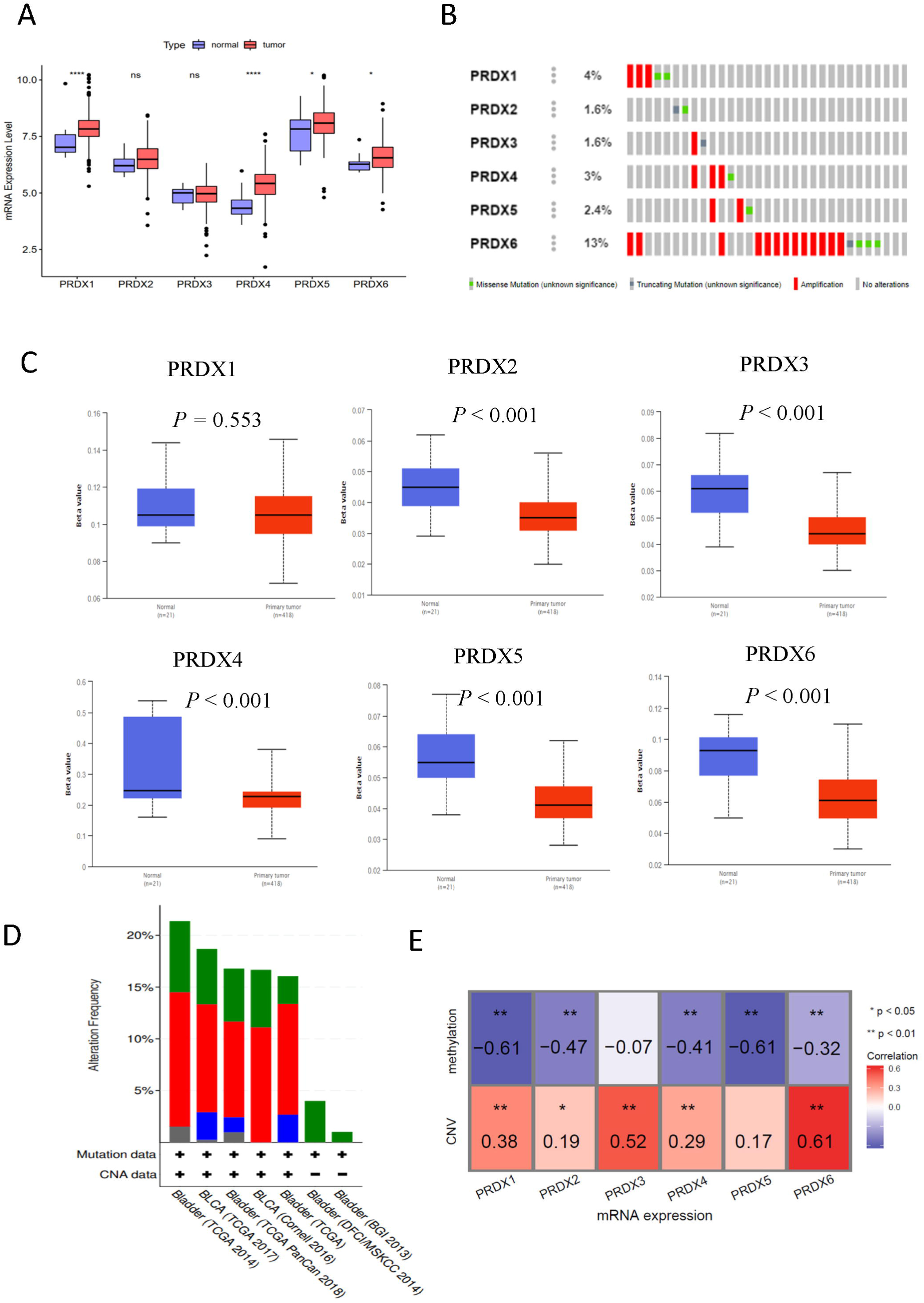
Expression profiles and genetic alterations of PRDXs in bladder cancer based on TCGA dataset. (A) The mRNA expression profiles of PRDXs in tumor compared to normal tissues. (B) The genetic alterations of PRDXs in bladder cancer. (C) The mRNA methylation levels of PRDXs in tumor compared to normal tissues. (D) The global percentage shows genetic alterations of PRDXs in different bladder cancer cohorts based on cBioprotal website. (E) Heatmap of the associations between PRDXs mRNA expression and CNV levels, DNA methylation in bladder cancer.

**Figure 5.**
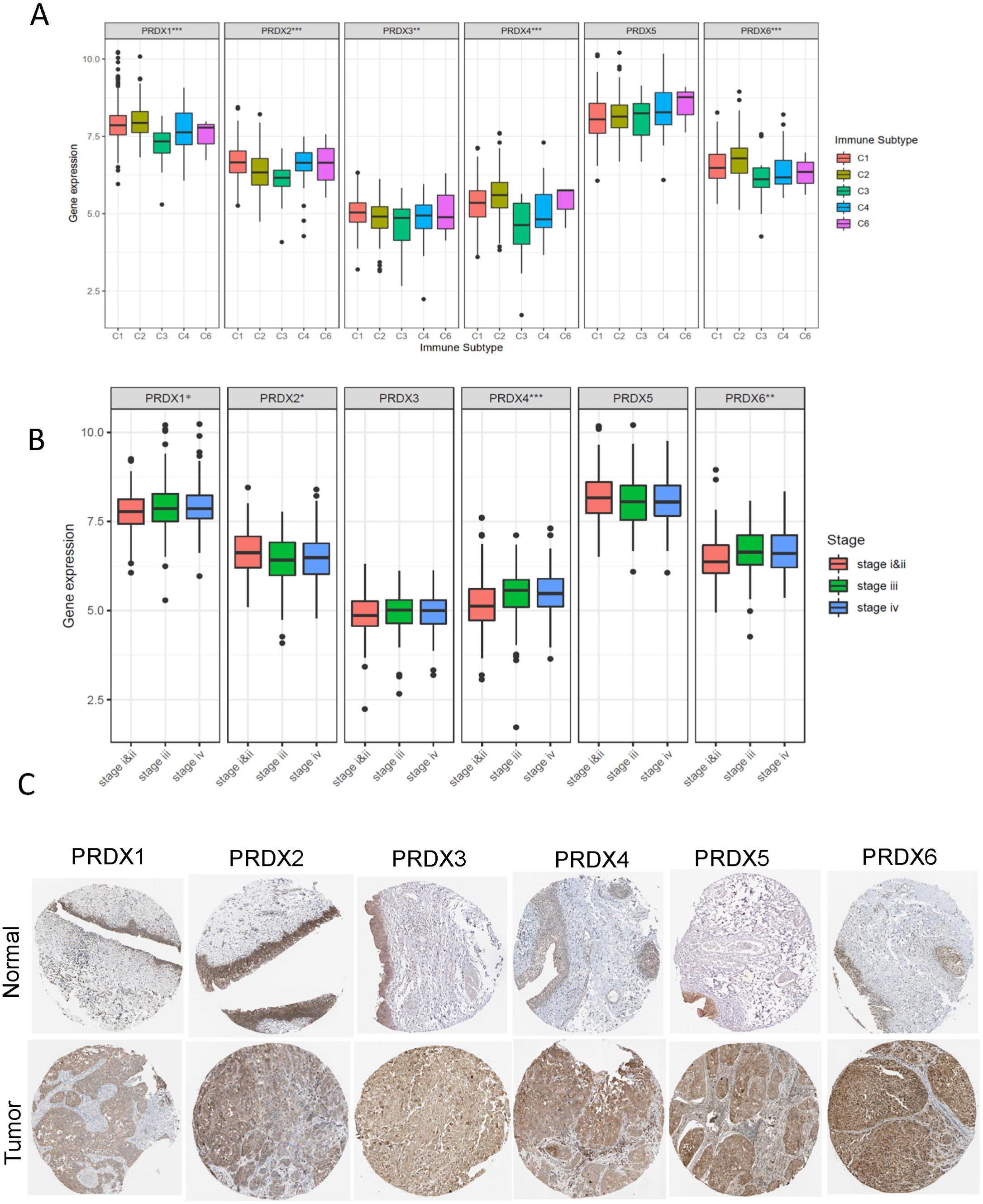
The PRDXs mRNA expression of different immune infiltrate subtypes, stage in bladder cancer and protein expression of PRDXs in tumor and normal bladder tissues. (A) Associations of PRDX gene expression with immune infiltrate subtypes. (B) Associations of PRDX gene expression with different stage. (C) Immunohistochemistry images obtained from the HPA database.

### The prognostic value and GSEA results of PRDXs in bladder cancer

To explore the prognostic value of PRDXs in LUAD, the Kaplan Meier (KM) plot with Log-rank tests was performed to determine the significance between PRDXs and OS. The results demonstrated that PRDX1 (*P* = 0.002), PRDX4 (*P* = 0.044) and PRDX6 (*P* = 0.02) were associated with poorer survival (**Figure 6A**). Subsequently, Gene set enrichment analysis (GSEA) methods was utilized to explore the potential biological pathways in BLCA of PRDXs between low and high expression data set. The significant pathways followed the criterion (FDR <0.05) in enrichment of MSigDB Collection (c2.cp.kegg.v7.0.symbols.gmt) were displayed in **Figure 6B**. The results demonstrated that PRDX1, PRDX2, PRDX4 and PRDX6 were associated with cell cycle, cell adhesion pathways (ECM receptor, focal adhesion, GAP junction, and cell adhesion molecules CAMs). PRDX1 were strongly associated with TGF-beta pathway, PRDX2 and PRDX6 with JAK-STAT pathway, PRDX3 with mTOR pathway. All PRDXs were closely with cell metabolism process, such as glycolysis gluconeogenesis, purine metabolism, fatty acid metabolism, drug metabolism cytochrome P450, retinol metabolism and metabolism of xenobiotics by cytochrome P450 (**Figure 6B**).

**Figure 6.**
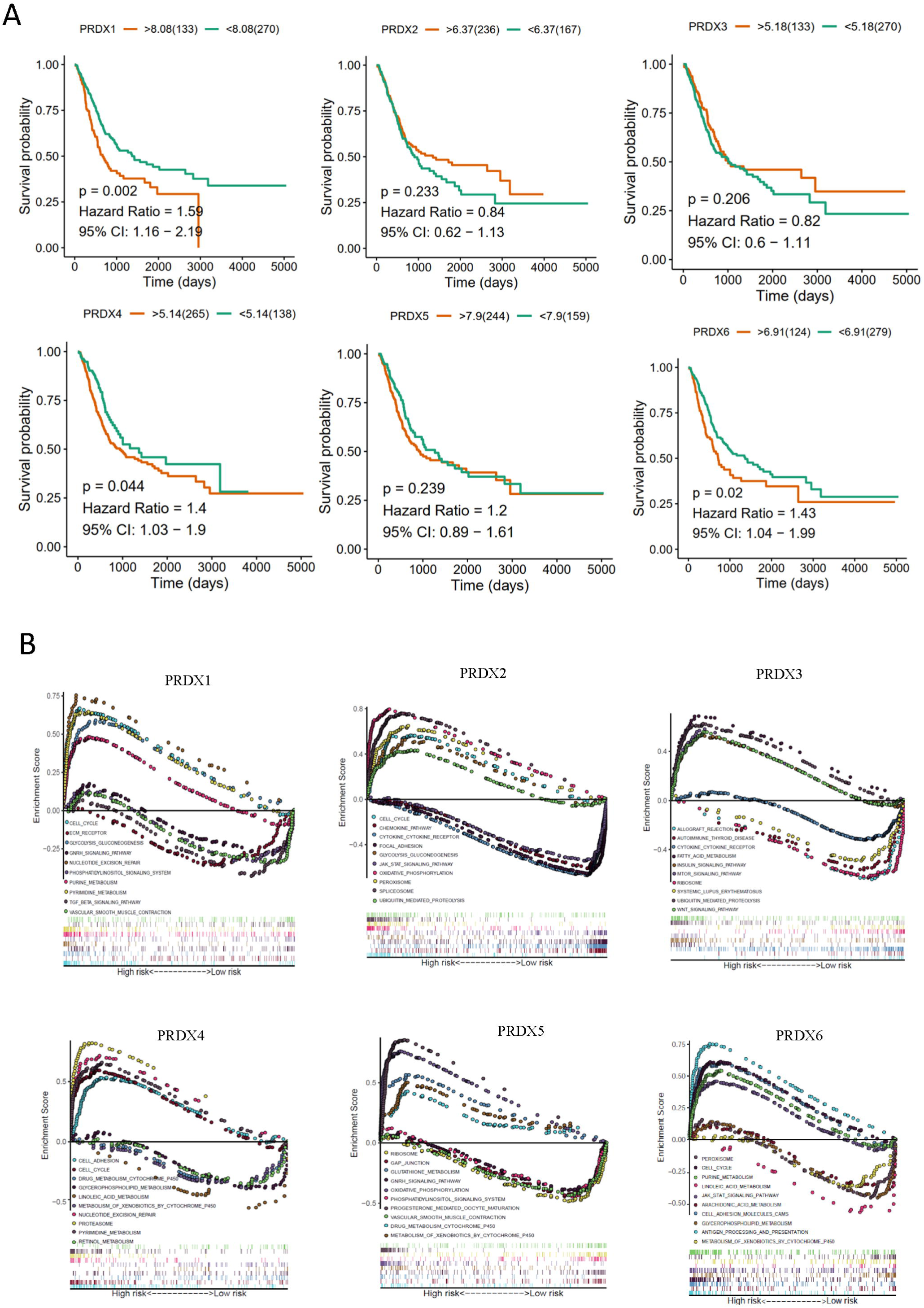
The overall survival and GSEA plots of RPDXs in bladder cancer based on TCGA. (A) The overall survival curves of PRDX genes. (B) The KEGG enrichment results of PRDX genes using GSEA method.

### Knockdown of PRDX6 inhibits the cell proliferation of BLCA

In order to verify the important role of PRDXs in bladder cancer, we selected PRDX6 which had high expression in tumor tissues with poor prognosis, and high genetic alterations rate in BLCA. We first knocked down PRDX6 in T24 and TCCSUP cell lines (**Figure 7A**), then tested if PRDX6 had any biological functions in BLCA cells. To examine the effects of PRDX6 on cell proliferation in BLCA cells, we firstly used CCK8 assay to detect the growth, the results revealed that knockdown PRDX6 in T24 and TCCSUP cells considerably suppressed the cell growth (**Figure 7B**). Similarly, colony formation assay also confirmed that knockdown of PRDX6 in T24 and TCCSUP cells reduced the colony number compared to the corresponding controls (**Figure 7C-D**). Next, we used FACS analysis to investigate how PRDX6 modulates cell cycle progression. We found that PRDX6 deletion resulted in an increased percentage of G2/M phase and a decreased percentage of cells in the S phase cell population both in T24 and TCCSUP cells (**Figure 7E-F**). What’s more, to investigate the effects of PRDX6 on apoptosis, the Annexin V-APC and 7-AAD staining was performed to assess. The scatter plots demonstrated a higher apoptotic index both in the T24 and TCCSUP cells with shPRDX6 (**Figure 7G-H**).

**Figure 7.**
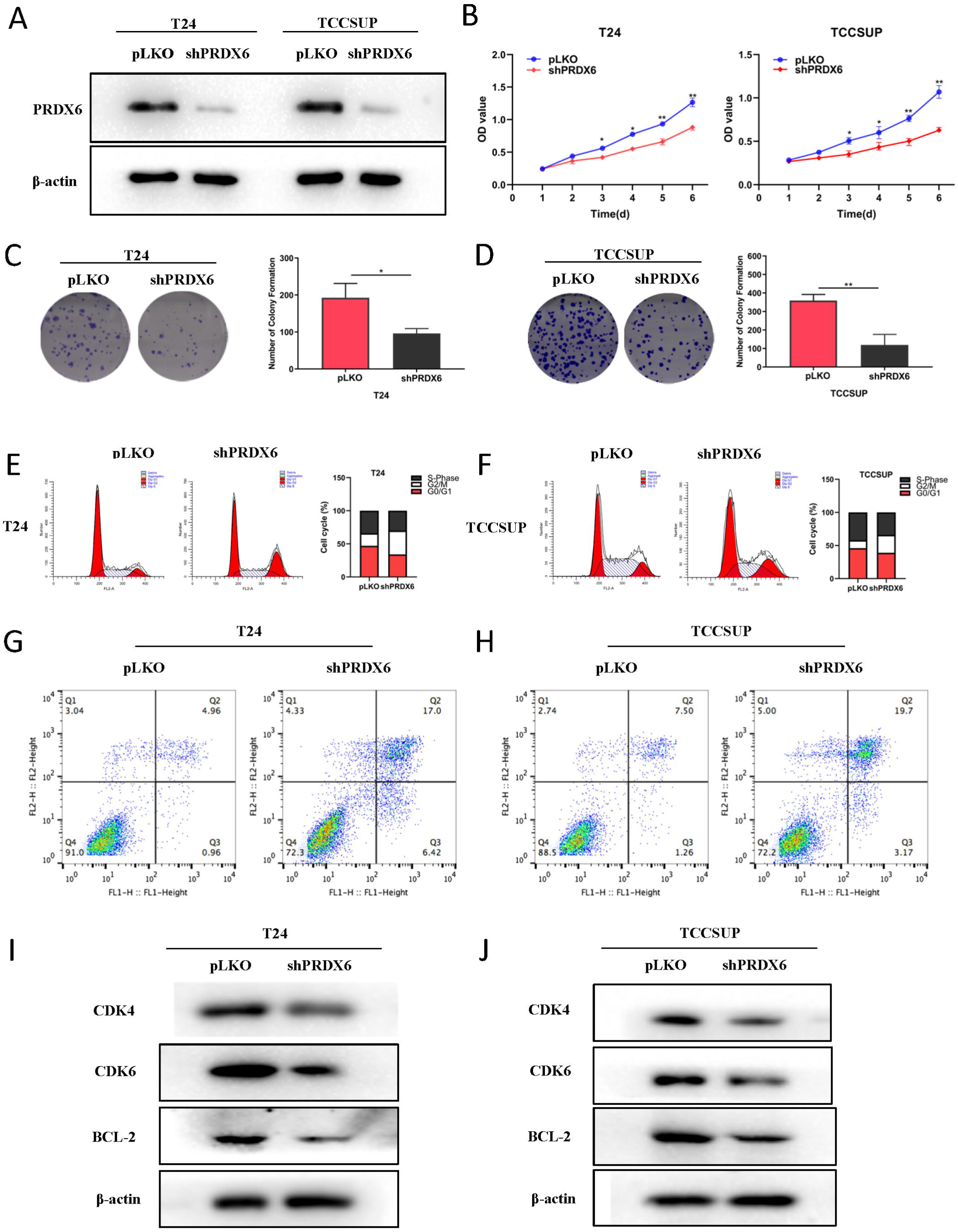
Knockdown of PRDX6 inhibits the cell proliferation of BLCA. (A) Knockdown efficiency of PRDX6 in T24 cells and TCCSUP cells. (B) CCK8 assay revealed cell proliferation in T24 cells and TCCSUP cells. (C-D) PRDX6 depletion decreased colony forming ability of T24 cells(C) and TCCSUP cells(D). (E-F) Knockdown PRDX6 arrested cell population in G2/M-phase both in T24 cells(E) and TCCSUP cells(F) by flow cytometer. (G-H) Knockdown PRDX6 increase the apoptosis of T24 cells(G) and TCCSUP cells(H). (I-J) Knockdown PRDX6 decrease the protein level of CDK4, CDK6 and BCL2. \**P* < 0.05, \*\**P* < 0.01.

Our previous analysis suggested that PRDX6 was highly correlated with the cell cycle. After knocking down PRDX6, we detected CDK4, CDK6, and BCL-2 proteins. The result showed that knockdown PRDX6 can significantly decrease the protein levels of CDK4, CDK6 and BCL-2 in the T24 cells (**Figure 7I**) and TCCSUP cells (**Figure 7J**). Collectively, these results demonstrated that knockdown of PRDX6 can inhibit the cell proliferation of BLCA.

### PRDX6 promotes bladder cancer cell proliferation via JAK2-STAT3 pathway

To investigate how PRDX6 regulated cell proliferation signaling, the GSEA analysis and found JAK-STAT3 pathway was significantly enriched in PRDX high group (**Figure 6E, Figure 8A**). STAT3 plays as an important role in urological related tumors^34^, and previous study has confirmed that STAT3 as the upstream signaling of CDK4/6 pathway^35^. Taken together, we speculated that JAK-STAT3 signaling can be regulated by PRDX6 to impact the proliferation of BLCA cells. As observed in **Figure 8B-C**, we utilized the WB to detect and found that PRDX6 knockdown decreased the JAK2 protein level, phosphorylation and total STAT3 protein level both in T24 and TCCSUP cells. Next, the results showed that the decreased total STAT3, CDK4, CDK6 and BCL-2 by shPRDX6 can be reversed under the addition of oeSTAT3 (**Figure 8D-E**). What’s more, the result of CCK8 (**Figure 8F-G**) and colony formation (**Figure 8H-K**) demonstrated that the diminished effect of proliferation by knockdown PRDX6 also can be reverse by oeSTAT3 in T24 and TCCSUP cells. Thus, these data suggested that PRDX6 regulates JAK2-STAT3 signaling, which probably contributes to promote the cell proliferation in BLCA.

**Figure 8.**
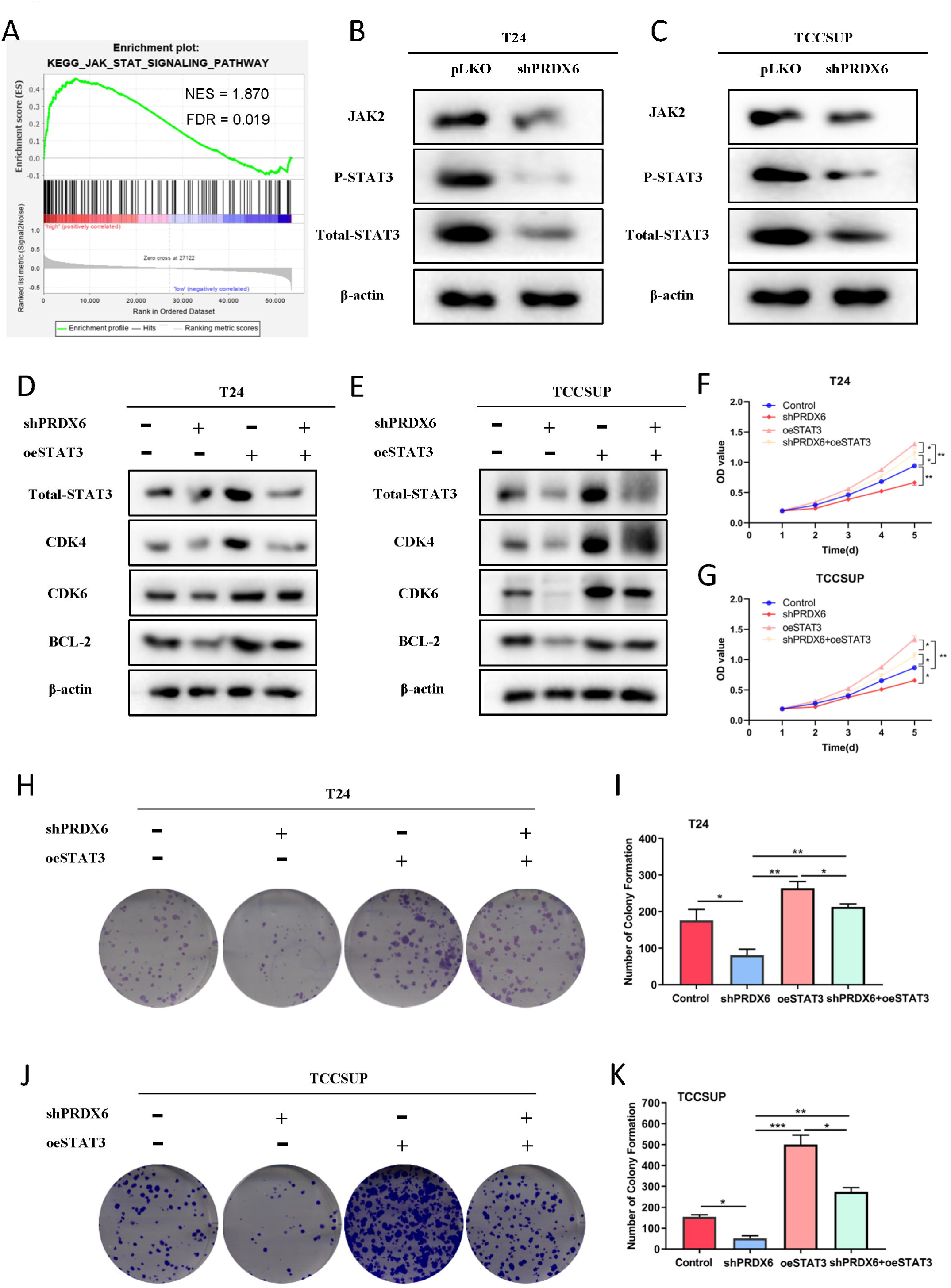
PRDX6 promotes bladder cancer cell proliferation via JAK2-STAT3 pathway. (A)The GSEA plot of JAK/STAT pathway between PRDX6 high and low group. (B-C) Western blot was used to test JAK2, p-STAT3 and STAT3 protein both in T24 cells(B) and TCCSUP cells(C). (D-G) To test whether PRDX6 regulate the cell growth via JAK2-STAT3 pathway, western blot assays and CCK8 were performed on T24 cells(D,F) and TCCSUP cells(E,G) transfected with control, shPRDX6, oeSTAT3, shPRDX6+oeSTAT3. (H-K) Colony formation were performed on T24 cells(H,I) and TCCSUP cells(J,K) transfected with control, shPRDX6, oeSTAT3, shPRDX6+oeSTAT3. *P < 0.05, **P < 0.01.

## Discussion

Accumulating researches demonstrated that PRDXs played vital roles in the progression of cancers, however, a systematic analyse of PRDXs in different cancers is deficiency. Therefore, it is necessary to study the expression, genetic alterations, regulation patterns and potential drugs for diagnosis and treatment of cancers with aberrantly PRDXs expression. The aim of our study was to investigate the characteristic features of PRDXs in pan-cancer and further explore the potential roles in BLCA. We found that the distributions of PRDXs mRNA expression levels among 33 tumor types were great different. PRDX1, PRDX2, PRDX4 and PRDX5 were mainly up-regulated, while PRDX3 and PRDX6 were primarily down-regulated expressions in several cancers. However, only PRDX4, PRDX6 were associated with the OS of patients in more than 8 cancer types, the rest PRDX members in less than 7 types. As we all known, epigenetic aberrations involve the initiation and progression of tumors^36^. Next, the genetic alterations and methylation of PRDXs were explored, which showed that PRDX gene expressions were positively correlated with CNV and negatively with methylation, which suggested that PRDXs probably are epigenetic-drive genes.

Subsequently, the correlations between PRDXs and immune subtypes, tumor stemness demonstrated that PRDX1, PRDX4, PRDX6 were closely related with type C1, C2 and C6 infiltrates, while tumor stemness scores were positively with all PRDX members, which indicated poor prognosis in pan-cancer. Considering that tumor stemness were strongly related with drug response, then the relationships between 10 tumor-related pathways and potential drugs were investigated. The results showed that PRDXs were associated with apoptosis and cell cycle pathway. By using PharmacoDB data, we found that increased expression of PRDX1, PRDX3, PRDX6 were closely related with a number of FDA approved EGFR/VEGFR inhibitor drugs, such as Pazopanib, Vandetanib, Lapatinib and Cediranib (*p*<0.001), which are used to treat tumors such as renal cell carcinoma^37^, melanoma, thyroid cancer^38^. This suggests that reasonable use of EGF/VEGF inhibitors can effectively treat some cancers with up-regulated expression of PRDXs. Besides, Gemcitabine, Doxorubicin, AZD8055 (mTOR inhibitor), GDC-0941 (PI3K inhibitor), Irinotecan (cytotoxic chemotherapy drug), 17-AAG (HSP90 inhibitor), Mitomycin-C (cytotoxic), and temsirolimus drugs were also strongly related with high expression of PRDXs, which control cancer by changing cancer metabolism. Genetic alterations and methylation results demonstrated that PRDX members were significantly positive correlated with CNV and methylation, which indicates that the combination of epigenetic inhibitors and EGF/VEGF inhibitors may be an effective anticancer treatment.

Considering that it had been reported PRDX expressions were aberrant and associated with OS, but most only focused on several members, systematic analyze of PRDXs in BLCA is needed. Therefore, in our study the characteristics of PRDXs in BLCA were explored. The results demonstrated that PRDX1, PRDX4, PRDX5 and PRDX6 were significant difference between tumor and normal tissues, which were also validated in HPA database. The KM plot showed the high expressions of PRDX1, PRDX4 and PRDX6 were associated with poor prognosis, which were consistent with the researches of Soini et al^39^ and Quan et al^30^. Finally, we chose the PRDX6, which has 13% genetic alterations and the correlations with CNV, methylation was 0.61 (*P* <0.01) and −0.32 (*P* < 0.01), to confirm the functions of PRDXs in BLCA. The T24 and TCCSUP bladder cancer cell lines were used to knockdown PRDX6 and test cell growth and cell cycle. Our results showed that the decreased expression of PRDX6 could inhibit the growth of cancer cells, which were in accord with the outcomes of esophageal carcinoma^40^, breast cancer^26^. The GSEA results showed that PRDX6 were positively associated with JAK/STAT pathway, then we detect the famous modules of JAK/STAT, JAK2, STAT3 and p-STAT3. The results exhibited that PRDX6 could regulate the cell proliferation via JAK2-STAT3 pathway and involve the process of cell cycle, which were agreed with the results in lung cancer model^41^ and may provide novel target for the treatment of BLCA.

In summary, our study provides systematic analyses of the characteristics of the PRDX gene family from expression profiles, associations with overall survival, genetic alterations, methylation, potential biological pathways and drugs in pan-cancer. Moreover, we comprehensively analyze the roles of PRDXs in BLCA and validate that PRDX6 could regulate cell proliferation via JAK2-STAT3 signaling and involve the cell cycle process, which may be more valuable for personalized treatment of cancer. In a word, we not only uncover the pivotal roles of the PRDXs in the progression of 33 types cancers, but validate the PRDX6 could attenuate the cell growth through JAK2-STAT3 pathway in BLCA.

## Materials and methods

### TCGA pan-cancer data

The 33 cancer types data, including RNA-seq, clinical information, stemness scores based on mRNA (mRNAsi) and immune subtypes were obtained from Xena browser (https://xenabrowser.net/datapages/). Among them, only 18 cancer types have adjacent normal tissue samples, which were used to investigate whether there was aberrantly gene expression in tumors compared to normal tissues. In additional, in order to analyze the correlation between gene expression (as continuous variable) and patient prognosis, patients lack of OS information were excluded.

### Tumor immune subtypes analysis

Six immune subtypes were classed to evaluate immune infiltrates situations in tumor environment^42^. Immune subtype was used to detect the relationship between PRDXs expression and immune infiltrate types in tumor microenvironment using ANOVA methods. Tumor stemness data obtained from TCGA dataset were performed to assess the stem-cell-like levels of tumor cells^32^. The association between cancer stemness and PRDXs expression profiles was examined using Spearman correlation method.

### Genetic alteration and DNA methylation

Genetic alterations, including mutation and copy number variation (CNV) of different cancer types were obtained from cBioPortal (http://www.cbioportal.org/). The association of CNV, methylation and tumor types were analyzed by GSCALite (http://bioinfo.life.hust.edu.cn/web/GSCALite/). The correlation between mRNA expression levels of PRDXs and CNV levels, methylation in BLCA were released from cBioPortal. The methylation levels between tumor and normal tissues in BLCA were obtained from UALCAN (http://ualcan.path.uab.edu).

### Drug and pathway analysis

The global proportion plot and heatmap of the PRDXs genes in 10 tumor-assocated pathways were acquired from GSCALite (http://bioinfo.life.hust.edu.cn/web/GSCALite/). All results for PRDXs-related drugs were obtained from PharmacoDB (https://pharmacodb.pmgenomics.ca/), and the meaningful cut-off is absolute value of correlation more than 0.1 and p<0.05 (Supplementary Table 3).

### Verification of expression profiles and pathway analyses of PRDXs

The Human Protein Atlas (HPA) database (http://www.proteinatlas.org/) was performed to verify the protein level of PRDXs. The prognostic value of the PRDXs was verified by using survival (https://github.com/therneau/survival) and survimer (https://github.com/kassambara/survminer/) package. GSEA(http://software.broadinstitute.org/gsea/index.jsp) package was applied to analyze the differential pathways based on KEGG gene sets.

### Cell culture and lentiviral of PRDX6, STAT3

T24, TCCSUP and HEK293T cell lines were purchased from the American Type Culture Collection company (ATCC, Manassas, VA), which were cultured in DMEM media and maintained in a humidified 5% CO2 environment at 37 °C^43^. Lentiviral pLKO-vec, pLKO-shPRDX6 were transfected into 293T cells and harvested according to our previous paper ^44^. For shPRDX6 knocked-down stable expression, puromycin (2μg/ml) was added to select stably transduced cells. The lentiviral soups were stored in −80 °C for later use. The constitutively active oeSTAT3 lentiviral vector (plasmid #24983) was obtained from Addgene (Cambridge, MA, USA).

### Western blot assay

Cells were lysed, quantified and equal 30µg protein, was separated on 10% SDS/PAGE gel, and then transferred onto PVDF membranes (Millipore, Billerica, MA). After blocked for 2 hours with 5% BSA at room temperature, the membrane was incubated in primary antibodies at 4°C for overnight. Next, the membrane was rinsed 3 times and incubated with secondary antibodies for 1 hour at room temperature. Finally, the protein bands imaged by ECL system (Thermo Fisher Scientific). The primary antibodies used in the study for western blot were listed below: β-actin (Santa Cruz, #sc-8432, CA), PRDX6 (Abcam, #ab59543, USA), CDK4 (Santa Cruz, #sc-56277, CA), CDK6 (Santa Cruz, #sc-39049, CA), BCL-2 (Santa Cruz, #sc-20067, CA), JAK2 (Santa Cruz, #sc-390539, CA), p-STAT3 (Santa Cruz, #sc-8059, CA), STAT3 (Santa Cruz, #sc-8019, CA).

### Cell viability assay, Colony Formation Assay and cell cycle analysis

Stably expressed cells (5×10^3^/well) were seeded in 96-well plates. Cell viability was measured with a Cell Counting Kit-8 (CCK-8). Cells were plated into 6-well plates at a density of 1×10^3^ cells per well. After 14 days. colonies were rinsed with PBS twice, fixed with 100% methanol, and dyeing with 0.1% crystal violet, then the cell numbers were calculated. For cell cycle assay, cells were harvested and washed with cold PBS, and fixed in 70% cold ethanol. The cells were stained with 50 mg/L propidium iodide for 30 min, before analysis with a fluorescence-activated cell sorter (BD FACS Flow Cytometer).

### Statistical analysis

The R-3.6.2 software was performed to analyze statistics and generate images. Comparison of PRDXs mRNA expression between tumor and normal tissues were performed using Wilcox test. The comparison of multi-groups was used ANOVA methods. Spearman’s correlation analysis was utilized to explore the correlation between continuous variables. Univariate cox proportional hazard regression models or Log-rank tests were used to assess the correlations between gene expression and patient overall survival. *P* vale < 0.05 was set as the significant different. Each statistical test was double-sided.

## Acknowledgments

This work was supported by Young Scientists Foundation of Changzhou No. 2 People’s Hospital (2018K009), High-Level Medical Talents Training Project NO.2016CZBJ035) and Changzhou Science and Technology Project (CJ20190100).

## Author Contributions

L.G. and J.L.M conceived and designed the project, L.G, J.L.M, C.Y, X.W, Q.S, H.W performed the experiments, analyzed the data. L.G, J.L.M and ZZ wrote the manuscript. S.L.G, S.F and LZ collected and analyzed the data and critically viewed and supervised the study.

## References

1. Nelson, KJ, Knutson, ST, Soito, L, Klomsiri, C, Poole, LB, and Fetrow, JS (2011). Analysis of the peroxiredoxin family: using active-site structure and sequence information for global classification and residue analysis. Proteins 79: 947–964.

2. Neumann, CA, and Fang, Q (2007). Are peroxiredoxins tumor suppressors? Curr Opin Pharmacol 7: 375–380.

3. Lee, YJ (2020). Knockout Mouse Models for Peroxiredoxins. Antioxidants (Basel) 9.

4. Rhee, SG, Kang, SW, Chang, TS, Jeong, W, and Kim, K (2001). Peroxiredoxin, a novel family of peroxidases. IUBMB Life 52: 35–41.

5. Hall, A, Karplus, PA, and Poole, LB (2009). Typical 2-Cys peroxiredoxins--structures, mechanisms and functions. FEBS J 276: 2469–2477.

6. Soito, L, Williamson, C, Knutson, ST, Fetrow, JS, Poole, LB, and Nelson, KJ (2011). PREX: PeroxiRedoxin classification indEX, a database of subfamily assignments across the diverse peroxiredoxin family. Nucleic Acids Res 39: D332–337.

7. Kumar, Y, Biswas, T, Thacker, G, Kanaujiya, JK, Kumar, S, Shukla, A, Khan, K, Sanyal, S, Chattopadhyay, N, Bandyopadhyay, A, et al. (2018). BMP signaling-driven osteogenesis is critically dependent on Prdx-1 expression-mediated maintenance of chondrocyte prehypetrophy. Free Radic Biol Med 118: 1–12.

8. Huh, JY, Kim, Y, Jeong, J, Park, J, Kim, I, Huh, KH, Kim, YS, Woo, HA, Rhee, SG, Lee, KJ, et al. (2012). Peroxiredoxin 3 is a key molecule regulating adipocyte oxidative stress, mitochondrial biogenesis, and adipokine expression. Antioxid Redox Signal 16: 229–243.

9. Jeong, SJ, Kim, S, Park, JG, Jung, IH, Lee, MN, Jeon, S, Kweon, HY, Yu, DY, Lee, SH, Jang, Y, et al. (2018). Prdx1 (peroxiredoxin 1) deficiency reduces cholesterol efflux via impaired macrophage lipophagic flux. Autophagy 14: 120–133.

10. Kim, SU, Park, YH, Kim, JM, Sun, HN, Song, IS, Huang, SM, Lee, SH, Chae, JI, Hong, S, Sik Choi, S. et al. (2014). Dominant role of peroxiredoxin/JNK axis in stemness regulation during neurogenesis from embryonic stem cells. Stem Cells 32: 998–1011.

11. Fujii, J, and Ikeda, Y (2002). Advances in our understanding of peroxiredoxin, a multifunctional, mammalian redox protein. Redox Rep 7: 123–130.

12. Wang, X, Phelan, SA, Petros, C, Taylor, EF, Ledinski, G, Jurgens, G, Forsman-Semb, K, and Paigen, B (2004). Peroxiredoxin 6 deficiency and atherosclerosis susceptibility in mice: significance of genetic background for assessing atherosclerosis. Atherosclerosis 177: 61–70.

13. Lv, WP, Li, MX, and Wang, L (2017). Peroxiredoxin 1 inhibits lipopolysaccharide-induced oxidative stress in lung tissue by regulating P38/JNK signaling pathway. Eur Rev Med Pharmacol Sci 21: 1876–1883.

14. Ding, C, Fan, X, and Wu, G (2017). Peroxiredoxin 1 – an antioxidant enzyme in cancer. J Cell Mol Med 21: 193–202.

15. Kim, JH, Lee, JM, Lee, HN, Kim, EK, Ha, B, Ahn, SM, Jang, HH, and Lee, SY (2012). RNA-binding properties and RNA chaperone activity of human peroxiredoxin 1. Biochem Biophys Res Commun 425: 730–734.

16. Lee, DJ, Kang, DH, Choi, M, Choi, YJ, Lee, JY, Park, JH, Park, YJ, Lee, KW, and Kang, SW (2013). Peroxiredoxin-2 represses melanoma metastasis by increasing E-Cadherin/beta-Catenin complexes in adherens junctions. Cancer Res 73: 4744–4757.17.

17. Hu, JX, Gao, Q, and Li, L (2013). Peroxiredoxin 3 is a novel marker for cell proliferation in cervical cancer. Biomed Rep 1: 228–230.

18. Kim, YS, Lee, HL, Lee, KB, Park, JH, Chung, WY, Lee, KS, Sheen, SS, Park, KJ, and Hwang, SC (2011). Nuclear factor E2-related factor 2 dependent overexpression of sulfiredoxin and peroxiredoxin III in human lung cancer. Korean J Intern Med 26: 304–313.

19. Choi, JH, Kim, TN, Kim, S, Baek, SH, Kim, JH, Lee, SR, and Kim, JR (2002). Overexpression of mitochondrial thioredoxin reductase and peroxiredoxin III in hepatocellular carcinomas. Anticancer Res 22: 3331–3335.

20. Karihtala, P, Mantyniemi, A, Kang, SW, Kinnula, VL, and Soini, Y (2003). Peroxiredoxins in breast carcinoma. Clin Cancer Res 9: 3418–3424.

21. Jiang, H, Wu, L, Mishra, M, Chawsheen, HA, and Wei, Q (2014). Expression of peroxiredoxin 1 and 4 promotes human lung cancer malignancy. Am J Cancer Res 4: 445–460.

22. Chang, KP, Yu, JS, Chien, KY, Lee, CW, Liang, Y, Liao, CT, Yen, TC, Lee, LY, Huang, LL, Liu, SC, et al. (2011). Identification of PRDX4 and P4HA2 as metastasis-associated proteins in oral cavity squamous cell carcinoma by comparative tissue proteomics of microdissected specimens using iTRAQ technology. J Proteome Res 10: 4935–4947.

23. Pritchard, C, Mecham, B, Dumpit, R, Coleman, I, Bhattacharjee, M, Chen, Q, Sikes, RA, and Nelson, PS (2009). Conserved gene expression programs integrate mammalian prostate development and tumorigenesis. Cancer Res 69: 1739–1747.

24. Karihtala, P, Kauppila, S, Soini, Y, and Arja Jukkola, V (2011). Oxidative stress and counteracting mechanisms in hormone receptor positive, triple-negative and basal-like breast carcinomas. BMC Cancer 11: 262.

25. Pylvas, M, Puistola, U, Kauppila, S, Soini, Y, and Karihtala, P (2010). Oxidative stress-induced antioxidant enzyme expression is an early phenomenon in ovarian carcinogenesis. Eur J Cancer 46: 1661–1667.

26. Chang, XZ, Li, DQ, Hou, YF, Wu, J, Lu, JS, Di, GH, Jin, W, Ou, ZL, Shen, ZZ, and Shao, ZM (2007). Identification of the functional role of peroxiredoxin 6 in the progression of breast cancer. Breast Cancer Res 9: R76.

27. Raatikainen, S, Aaaltomaa, S, Karja, V, and Soini, Y (2015). Increased Peroxiredoxin 6 Expression Predicts Biochemical Recurrence in Prostate Cancer Patients After Radical Prostatectomy. Anticancer Res 35: 6465–6470.

28. Kalinina, EV, Berezov, TT, Shtil, AA, Chernov, NN, Glazunova, VA, Novichkova, MD, and Nurmuradov, NK (2012). Expression of peroxiredoxin 1, 2, 3, and 6 genes in cancer cells during drug resistance formation. Bull Exp Biol Med 153: 878–881.

29. Pak, JH, Choi, WH, Lee, HM, Joo, WD, Kim, JH, Kim, YT, Kim, YM, and Nam, JH (2011). Peroxiredoxin 6 overexpression attenuates cisplatin-induced apoptosis in human ovarian cancer cells. Cancer Invest 29: 21–28.

30. Quan, C, Cha, EJ, Lee, HL, Han, KH, Lee, KM, and Kim, WJ (2006). Enhanced expression of peroxiredoxin I and VI correlates with development, recurrence and progression of human bladder cancer. J Urol 175: 1512–1516.

31. Tamborero, D, Rubio-Perez, C, Muinos, F, Sabarinathan, R, Piulats, JM, Muntasell, A, Dienstmann, R, Lopez-Bigas, N, and Gonzalez-Perez, A (2018). A Pan-cancer Landscape of Interactions between Solid Tumors and Infiltrating Immune Cell Populations. Clin Cancer Res 24: 3717–3728.

32. Malta, TM, Sokolov, A, Gentles, AJ, Burzykowski, T, Poisson, L, Weinstein, JN, Kaminska, B, Huelsken, J, Omberg, L, Gevaert, O, et al. (2018). Machine Learning Identifies Stemness Features Associated with Oncogenic Dedifferentiation. Cell 173: 338–354 e315.

33. Smirnov, P, Kofia, V, Maru, A, Freeman, M, Ho, C, El-Hachem, N, Adam, GA, Ba-Alawi, W, Safikhani, Z, and Haibe-Kains, B (2018). PharmacoDB: an integrative database for mining in vitro anticancer drug screening studies. Nucleic Acids Res 46: D994–D1002.

34. Hindupur, SV, Schmid, SC, Koch, JA, Youssef, A, Baur, EM, Wang, D, Horn, T, Slotta-Huspenina, J, Gschwend, JE, Holm, PS, et al. (2020). STAT3/5 Inhibitors Suppress Proliferation in Bladder Cancer and Enhance Oncolytic Adenovirus Therapy. Int J Mol Sci 21.

35. Aboushousha, T, Hammam, O, Aref, A, Kamel, A, Badawy, M, and Abdel Hamid, A (2020). Tissue Profile of CDK4 and STAT3 as Possible Innovative Therapeutic Targets in Urinary Bladder Cancer. Asian Pac J Cancer Prev 21: 547–554.

36. Villanueva, L, Alvarez-Errico, D, and Esteller, M (2020). The Contribution of Epigenetics to Cancer Immunotherapy. Trends Immunol.

37. Li, W, Feng, C, Di, W, Hong, S, Chen, H, Ejaz, M, Yang, Y, and Xu, TR (2020). Clinical use of vascular endothelial growth factor receptor inhibitors for the treatment of renal cell carcinoma. Eur J Med Chem 200: 112482.

38. Lee, CS, Baek, J, and Han, SY (2017). The Role of Kinase Modulators in Cellular Senescence for Use in Cancer Treatment. Molecules 22.

39. Soini, Y, Haapasaari, KM, Vaarala, MH, Turpeenniemi-Hujanen, T, Karja, V, and Karihtala, P (2011). 8-hydroxydeguanosine and nitrotyrosine are prognostic factors in urinary bladder carcinoma. Int J Clin Exp Pathol 4: 267–275.

40. He, Y, Xu, W, Xiao, Y, Pan, L, Chen, G, Tang, Y, Zhou, J, Wu, J, Zhu, W, Zhang, S, et al. (2018). Overexpression of Peroxiredoxin 6 (PRDX6) Promotes the Aggressive Phenotypes of Esophageal Squamous Cell Carcinoma. J Cancer 9: 3939–3949.

41. Yun, HM, Park, KR, Park, MH, Kim, DH, Jo, MR, Kim, JY, Kim, EC, Yoon, DY, Han, SB, and Hong, JT (2015). PRDX6 promotes tumor development via the JAK2/STAT3 pathway in a urethane-induced lung tumor model. Free Radic Biol Med 80: 136–144.

42. Thorsson, V, Gibbs, DL, Brown, SD, Wolf, D, Bortone, DS, Ou Yang, TH, Porta-Pardo, E, Gao, GF, Plaisier, CL, Eddy, JA, et al. (2018). The Immune Landscape of Cancer. Immunity 48: 812–830 e814.

43. Gao, L, Meng, J, Zhang, M, Fan, S, Gao, S, Wang, X, and Liang, C (2020). Expression and Prognostic Values of the Roof Plate-Specific Spondin Family in Bladder Cancer. DNA Cell Biol 39: 1072–1089.

44. Gu, J, Zhang, Y, Han, Z, Gao, L, Cui, J, Sun, Y, Niu, Y, You, B, Huang, CP, Chang, C, et al. (2020). Targeting the ERbeta/Angiopoietin-2/Tie-2 signaling-mediated angiogenesis with the FDA-approved anti-estrogen Faslodex to increase the Sunitinib sensitivity in RCC. Cell Death Dis 11: 367.

