## Supplemental Table 1 for "Integrative Analysis the characterization of peroxiredoxins in pan-cancer"

| cancer | PRDX1 | | | PRDX2 | | | PRDX3 | | |
| --- | --- | --- | --- | --- | --- | --- | --- | --- | --- |
|  | HR | 95%CI | pvalue | HR | 95%CI | pvalue | HR | 95%CI | pvalue |
| ACC | 1.42 | 0.8-2.52 | 0.23 | 1.12 | 0.66-1.9 | 0.67 | 0.63 | 0.37-1.07 | 0.09 |
| BLCA | 1.4 | 1.12-1.77 | <0.01 | 0.94 | 0.76-1.17 | 0.6 | 0.94 | 0.72-1.23 | 0.63 |
| BRCA | 1.18 | 0.92-1.51 | 0.2 | 1.07 | 0.82-1.39 | 0.62 | 1.12 | 0.84-1.47 | 0.44 |
| CESC | 0.97 | 0.68-1.39 | 0.87 | 0.66 | 0.51-0.86 | <0.01 | 1.16 | 0.79-1.7 | 0.46 |
| CHOL | 0.97 | 0.5-1.86 | 0.92 | 1.25 | 0.78-2.01 | 0.36 | 1.66 | 0.73-3.76 | 0.23 |
| COAD | 1.02 | 0.74-1.4 | 0.91 | 0.99 | 0.76-1.29 | 0.96 | 0.79 | 0.56-1.12 | 0.19 |
| DLBC | 0.25 | 0.06-1.06 | 0.06 | 1.24 | 0.69-2.25 | 0.47 | 0.77 | 0.19-3.12 | 0.72 |
| ESCA | 1.1 | 0.76-1.6 | 0.61 | 1.02 | 0.71-1.46 | 0.91 | 1.34 | 0.83-2.15 | 0.23 |
| GBM | 1.21 | 0.85-1.71 | 0.28 | 0.77 | 0.58-1.03 | 0.08 | 0.85 | 0.62-1.16 | 0.3 |
| HNSC | 1.13 | 0.91-1.4 | 0.26 | 0.9 | 0.79-1.04 | 0.15 | 1.3 | 1.01-1.68 | 0.05 |
| KICH | 1.46 | 0.28-7.73 | 0.65 | 0.94 | 0.23-3.93 | 0.94 | 1.45 | 0.51-4.14 | 0.49 |
| KIRC | 1.25 | 0.88-1.78 | 0.21 | 0.72 | 0.6-0.86 | <0.01 | 0.66 | 0.52-0.83 | <0.01 |
| KIRP | 1.6 | 1.06-2.41 | 0.02 | 0.51 | 0.38-0.69 | <0.01 | 0.95 | 0.59-1.52 | 0.83 |
| LAML | 1.37 | 1.01-1.86 | 0.04 | 1.01 | 0.86-1.19 | 0.87 | 1.2 | 0.75-1.92 | 0.44 |
| LGG | 1.25 | 1.04-1.5 | 0.02 | 0.49 | 0.31-0.79 | <0.01 | 0.34 | 0.21-0.56 | <0.01 |
| LIHC | 1.67 | 1.32-2.13 | <0.01 | 1 | 0.75-1.33 | 0.98 | 0.91 | 0.71-1.15 | 0.42 |
| LUAD | 1.08 | 0.91-1.29 | 0.39 | 1.05 | 0.88-1.26 | 0.56 | 1.08 | 0.84-1.39 | 0.54 |
| LUSC | 0.99 | 0.85-1.15 | 0.86 | 0.83 | 0.7-0.98 | 0.03 | 0.89 | 0.7-1.14 | 0.37 |
| MESO | 1.19 | 0.73-1.95 | 0.49 | 0.65 | 0.47-0.91 | 0.01 | 0.71 | 0.47-1.06 | 0.09 |
| OV | 0.89 | 0.71-1.11 | 0.3 | 0.94 | 0.79-1.12 | 0.51 | 0.86 | 0.69-1.07 | 0.19 |
| PAAD | 1.67 | 1.16-2.4 | 0.01 | 0.54 | 0.37-0.8 | <0.01 | 1.26 | 0.75-2.12 | 0.38 |
| PCPG | 0.72 | 0.22-2.36 | 0.59 | 2.17 | 0.34-13.89 | 0.41 | 0.6 | 0.17-2.1 | 0.42 |
| PRAD | 1.37 | 0.28-6.8 | 0.7 | 1.41 | 0.39-5.04 | 0.6 | 3.33 | 0.94-11.88 | 0.06 |
| READ | 0.64 | 0.25-1.63 | 0.35 | 0.72 | 0.34-1.5 | 0.37 | 1.17 | 0.55-2.47 | 0.69 |
| SARC | 1.12 | 0.84-1.49 | 0.45 | 0.95 | 0.81-1.12 | 0.56 | 0.64 | 0.43-0.94 | 0.02 |
| SKCM | 1.04 | 0.92-1.18 | 0.53 | 0.98 | 0.9-1.06 | 0.62 | 0.74 | 0.6-0.92 | 0.01 |
| STAD | 0.86 | 0.66-1.13 | 0.27 | 0.86 | 0.67-1.11 | 0.24 | 1.03 | 0.76-1.41 | 0.84 |
| TGCT | 1.22 | 0.36-4.07 | 0.75 | 0.72 | 0.3-1.72 | 0.45 | 1.02 | 0.19-5.45 | 0.98 |
| THCA | 0.54 | 0.19-1.54 | 0.25 | 0.56 | 0.18-1.7 | 0.31 | 0.84 | 0.25-2.79 | 0.77 |
| THYM | 1.66 | 0.41-6.66 | 0.47 | 1.54 | 0.45-5.29 | 0.49 | 1.3 | 0.38-4.41 | 0.67 |
| UCEC | 1.06 | 0.77-1.45 | 0.73 | 0.95 | 0.73-1.24 | 0.7 | 0.98 | 0.74-1.3 | 0.9 |
| UCS | 1.46 | 0.71-2.99 | 0.31 | 1.3 | 0.75-2.24 | 0.35 | 1.14 | 0.59-2.2 | 0.69 |
| UVM | 1.32 | 0.61-2.86 | 0.48 | 0.87 | 0.39-1.97 | 0.74 | 1.72 | 1.01-2.94 | 0.05 |

Table S1 The unicox results of PRDXs

(continue)

| cancer | PRDX4 | | | PRDX5 | | | PRDX6 | | |
| --- | --- | --- | --- | --- | --- | --- | --- | --- | --- |
|  | HR | 95%CI | pvalue | HR | 95%CI | pvalue | HR | 95%CI | pvalue |
| ACC | 0.85 | 0.61-1.18 | 0.34 | 0.88 | 0.54-1.44 | 0.62 | 2.24 | 1.18-4.28 | 0.01 |
| BLCA | 1.15 | 0.93-1.43 | 0.2 | 0.98 | 0.78-1.24 | 0.88 | 1.41 | 1.13-1.76 | <0.01 |
| BRCA | 1.19 | 0.95-1.49 | 0.14 | 0.99 | 0.8-1.23 | 0.93 | 1.18 | 0.86-1.61 | 0.3 |
| CESC | 1.04 | 0.71-1.54 | 0.82 | 0.91 | 0.65-1.28 | 0.59 | 1.05 | 0.68-1.63 | 0.82 |
| CHOL | 1.58 | 0.61-4.08 | 0.34 | 1.06 | 0.55-2.05 | 0.86 | 1.85 | 0.62-5.5 | 0.27 |
| COAD | 0.94 | 0.72-1.24 | 0.67 | 1.02 | 0.85-1.22 | 0.82 | 1.09 | 0.68-1.74 | 0.72 |
| DLBC | 0.68 | 0.2-2.35 | 0.54 | 1.09 | 0.44-2.67 | 0.85 | 0.95 | 0.43-2.11 | 0.9 |
| ESCA | 1.82 | 1.24-2.69 | <0.01 | 0.96 | 0.59-1.56 | 0.88 | 0.99 | 0.62-1.6 | 0.98 |
| GBM | 1.03 | 0.83-1.29 | 0.77 | 0.92 | 0.66-1.27 | 0.61 | 1.18 | 0.86-1.63 | 0.29 |
| HNSC | 1.21 | 0.98-1.49 | 0.08 | 1.13 | 0.92-1.37 | 0.23 | 1.41 | 1.11-1.8 | 0.01 |
| KICH | 4.29 | 1.72-10.72 | <0.01 | 1.25 | 0.3-5.24 | 0.76 | 6.49 | 0.91-46.19 | 0.06 |
| KIRC | 0.99 | 0.76-1.3 | 0.94 | 1.08 | 0.85-1.36 | 0.55 | 1.43 | 0.98-2.09 | 0.06 |
| KIRP | 4.03 | 2.36-6.9 | <0.01 | 0.55 | 0.32-0.95 | 0.03 | 2.39 | 1.43-3.99 | <0.01 |
| LAML | 1.57 | 1.19-2.09 | <0.01 | 2.01 | 1.35-2.99 | <0.01 | 1.46 | 0.72-2.94 | 0.29 |
| LGG | 1.52 | 1.15-2.01 | <0.01 | 0.47 | 0.29-0.77 | <0.01 | 1.38 | 1.14-1.68 | <0.01 |
| LIHC | 1.4 | 1.1-1.78 | 0.01 | 1.3 | 0.99-1.7 | 0.06 | 1.39 | 1.07-1.82 | 0.01 |
| LUAD | 1.07 | 0.9-1.26 | 0.45 | 0.99 | 0.84-1.17 | 0.92 | 1.24 | 0.96-1.6 | 0.1 |
| LUSC | 0.93 | 0.75-1.16 | 0.53 | 1.01 | 0.8-1.27 | 0.92 | 0.92 | 0.73-1.17 | 0.49 |
| MESO | 1.47 | 1.03-2.08 | 0.03 | 0.61 | 0.42-0.89 | 0.01 | 0.76 | 0.54-1.07 | 0.12 |
| OV | 0.83 | 0.69-1 | 0.05 | 0.74 | 0.6-0.91 | <0.01 | 0.71 | 0.53-0.96 | 0.03 |
| PAAD | 0.84 | 0.61-1.15 | 0.28 | 1.25 | 0.85-1.83 | 0.25 | 1.54 | 1-2.37 | 0.05 |
| PCPG | 1.37 | 0.62-3.05 | 0.43 | 2.04 | 0.52-8.03 | 0.31 | 2.26 | 0.7-7.31 | 0.17 |
| PRAD | 1.21 | 0.46-3.15 | 0.7 | 0.58 | 0.24-1.41 | 0.23 | 0.94 | 0.28-3.2 | 0.92 |
| READ | 1.04 | 0.56-1.93 | 0.9 | 1.08 | 0.69-1.67 | 0.74 | 1.65 | 0.75-3.6 | 0.21 |
| SARC | 1.21 | 0.98-1.49 | 0.08 | 0.95 | 0.74-1.22 | 0.7 | 0.96 | 0.79-1.16 | 0.67 |
| SKCM | 0.91 | 0.72-1.13 | 0.38 | 1.11 | 0.87-1.42 | 0.42 | 0.83 | 0.66-1.05 | 0.12 |
| STAD | 1.13 | 0.89-1.44 | 0.32 | 0.94 | 0.73-1.22 | 0.65 | 1.18 | 0.81-1.71 | 0.39 |
| TGCT | 1.29 | 0.25-6.64 | 0.76 | 0.91 | 0.14-6.09 | 0.92 | 0.54 | 0.13-2.19 | 0.39 |
| THCA | 0.89 | 0.35-2.29 | 0.81 | 0.59 | 0.24-1.4 | 0.23 | 0.81 | 0.18-3.64 | 0.78 |
| THYM | 1.22 | 0.41-3.61 | 0.72 | 1.18 | 0.45-3.11 | 0.74 | 5.09 | 1.18-21.86 | 0.03 |
| UCEC | 0.94 | 0.73-1.21 | 0.63 | 1.06 | 0.79-1.42 | 0.68 | 1.64 | 1.16-2.31 | <0.01 |
| UCS | 1.06 | 0.66-1.71 | 0.8 | 0.81 | 0.44-1.47 | 0.48 | 0.92 | 0.55-1.53 | 0.75 |
| UVM | 3.63 | 1.69-7.82 | <0.01 | 3.41 | 1.46-7.98 | <0.01 | 0.35 | 0.07-1.7 | 0.19 |
