## Supplemental Table 2 for "Integrative Analysis the characterization of peroxiredoxins in pan-cancer"

Table S2 The associations of PRDXs and mRNAsi.

| CancerType | PRDX1 | | PRDX2 | | PRDX3 | |
| --- | --- | --- | --- | --- | --- | --- |
|  | cor | pvalue | cor | pvalue | cor | pvalue |
| TGCT | 0.662515 | 0 | -0.06596 | 0.414386 | 0.26808 | 0.000775 |
| UCEC | 0.499293 | 0 | 0.320639 | 6.16E-14 | 0.246003 | 1.18E-08 |
| BRCA | 0.456143 | 0 | 0.236001 | 3.99E-15 | 0.062662 | 0.038969 |
| HNSC | 0.447386 | 0 | 0.454016 | 0 | 0.190253 | 2.07E-05 |
| LUSC | 0.432584 | 0 | 0.458464 | 0 | 0.205984 | 4.91E-06 |
| UCS | 0.431032 | 0.001013 | 0.459945 | 0.000418 | 0.225632 | 0.094539 |
| LUAD | 0.417209 | 0 | 0.214372 | 1.12E-06 | 0.460025 | 0 |
| COAD | 0.390515 | 0 | 0.357645 | 1.13E-14 | 0.416368 | 0 |
| KIRP | 0.388799 | 1.55E-11 | 0.271295 | 3.82E-06 | 0.408927 | 8.98E-13 |
| OV | 0.382456 | 1.76E-10 | 0.351494 | 5.47E-09 | 0.368605 | 8.60E-10 |
| LIHC | 0.368428 | 3.32E-13 | 0.153647 | 0.003074 | -0.04513 | 0.386505 |
| STAD | 0.363988 | 1.21E-12 | 0.383337 | 4.92E-14 | 0.385884 | 3.07E-14 |
| SARC | 0.315967 | 2.53E-07 | 0.125838 | 0.043493 | 0.186771 | 0.002633 |
| READ | 0.309497 | 9.13E-05 | 0.305394 | 0.000114 | 0.3868 | 7.54E-07 |
| CESC | 0.302641 | 9.00E-08 | 0.279571 | 8.46E-07 | 0.013018 | 0.821357 |
| MESO | 0.287042 | 0.007555 | -0.05522 | 0.612938 | 0.232228 | 0.031648 |
| BLCA | 0.276518 | 1.83E-08 | 0.404675 | 0 | 0.166558 | 0.000789 |
| KIRC | 0.267753 | 8.05E-10 | 0.214567 | 9.68E-07 | 0.363213 | 8.25E-18 |
| PAAD | 0.265948 | 0.000823 | 0.263264 | 0.000933 | 0.125365 | 0.118835 |
| ESCA | 0.257846 | 0.001031 | 0.356243 | 4.39E-06 | 0.448271 | 4.11E-09 |
| CHOL | 0.250965 | 0.139583 | 0.098842 | 0.564956 | 0.198713 | 0.244315 |
| GBM | 0.216541 | 0.005156 | 0.484728 | 4.26E-11 | 0.00322 | 0.967132 |
| PRAD | 0.192063 | 1.86E-05 | 0.345228 | 4.19E-15 | 0.433857 | 0 |
| THCA | 0.171614 | 0.000109 | 0.332477 | 2.28E-14 | 0.394367 | 0 |
| SKCM | 0.165132 | 0.000325 | -0.05564 | 0.228042 | 0.25464 | 2.35E-08 |
| LAML | 0.041478 | 0.619966 | 0.021973 | 0.792835 | 0.416194 | 2.52E-07 |
| DLBC | 0.03528 | 0.811404 | -0.01465 | 0.921231 | 0.310139 | 0.032367 |
| ACC | -0.08689 | 0.4486 | 0.475512 | 1.39E-05 | 0.227937 | 0.044946 |
| KICH | -0.10375 | 0.413622 | 0.498397 | 3.55E-05 | 0.078571 | 0.536229 |
| UVM | -0.14534 | 0.197928 | 0.071683 | 0.526709 | 0.215026 | 0.055594 |
| LGG | -0.40949 | 0 | 0.397624 | 0 | 0.19773 | 5.44E-06 |
| PCPG | -0.41279 | 9.69E-09 | 0.292609 | 6.57E-05 | 0.197628 | 0.007575 |
| THYM | -0.57007 | 0 | 0.666764 | 0 | 0.163018 | 0.076528 |

(continue)

| CancerType | PRDX4 | | PRDX5 | | PRDX6 | |
| --- | --- | --- | --- | --- | --- | --- |
|  | cor | pvalue | cor | pvalue | cor | pvalue |
| TGCT | 0.359035 | 5.21E-06 | -0.05548 | 0.492538 | 0.004921 | 0.951502 |
| UCEC | 0.273262 | 2.11E-10 | 0.041268 | 0.344275 | 0.36993 | 0 |
| BRCA | 0.461305 | 0 | 0.062224 | 0.040359 | 0.357279 | 0 |
| HNSC | 0.090357 | 0.044313 | 0.101004 | 0.024515 | 0.376125 | 0 |
| LUSC | 0.003724 | 0.934707 | 0.016492 | 0.716766 | 0.445858 | 0 |
| UCS | -0.01784 | 0.896022 | 0.241012 | 0.073706 | 0.085988 | 0.527581 |
| LUAD | 0.315692 | 4.05E-13 | 0.009767 | 0.825946 | 0.439484 | 0 |
| COAD | 0.313762 | 1.92E-11 | 0.144268 | 0.002383 | 0.26202 | 2.55E-08 |
| KIRP | 0.235281 | 6.53E-05 | 0.336008 | 7.89E-09 | 0.347797 | 2.18E-09 |
| OV | 0.273163 | 7.27E-06 | 0.211506 | 0.000556 | 0.280531 | 4.02E-06 |
| LIHC | 0.099751 | 0.055261 | 0.366381 | 4.62E-13 | 0.291637 | 1.28E-08 |
| STAD | 0.361655 | 1.74E-12 | 0.221967 | 2.13E-05 | 0.318356 | 7.13E-10 |
| SARC | 0.161406 | 0.009467 | 0.134931 | 0.030325 | 0.354139 | 6.22E-09 |
| READ | 0.175294 | 0.028729 | 0.127477 | 0.112713 | 0.245329 | 0.002077 |
| CESC | 0.129904 | 0.02379 | -0.01952 | 0.7349 | 0.274216 | 1.38E-06 |
| MESO | 0.057069 | 0.60108 | 0.276513 | 0.010163 | 0.338271 | 0.001529 |
| BLCA | 0.094718 | 0.057172 | -0.07398 | 0.13764 | 0.096825 | 0.05184 |
| KIRC | 0.168548 | 0.000126 | 0.2471 | 1.53E-08 | 0.327502 | 3.48E-14 |
| PAAD | 0.139856 | 0.081641 | 0.28816 | 0.00028 | 0.436598 | 1.73E-08 |
| ESCA | 0.293367 | 0.000177 | 0.187774 | 0.017536 | 0.236262 | 0.002693 |
| CHOL | 0.050193 | 0.770682 | 0.058172 | 0.735368 | 0.380952 | 0.022539 |
| GBM | -0.02262 | 0.772193 | 0.203355 | 0.008692 | -0.33106 | 1.48E-05 |
| PRAD | 0.383037 | 0 | -0.01697 | 0.70715 | 0.118887 | 0.008298 |
| THCA | -0.07716 | 0.083228 | 0.2027 | 4.61E-06 | 0.253386 | 8.71E-09 |
| SKCM | -0.07963 | 0.084297 | 0.090325 | 0.050128 | -0.02843 | 0.538035 |
| LAML | 0.369887 | 5.48E-06 | -0.27407 | 0.000886 | 0.334168 | 4.42E-05 |
| DLBC | -0.014 | 0.924734 | 0.052215 | 0.72377 | 0.348676 | 0.015583 |
| ACC | 0.099736 | 0.384186 | 0.25085 | 0.026996 | 0.204315 | 0.072854 |
| KICH | -0.01914 | 0.880476 | 0.071291 | 0.574741 | 0.262134 | 0.036689 |
| UVM | 0.227496 | 0.042618 | 0.023019 | 0.839105 | 0.105767 | 0.349711 |
| LGG | 0.039735 | 0.364468 | 0.275996 | 1.61E-10 | -0.55083 | 0 |
| PCPG | -0.2549 | 0.000536 | 0.237951 | 0.001253 | -0.07755 | 0.297788 |
| THYM | -0.38582 | 1.71E-05 | 0.043249 | 0.64003 | 0.296589 | 0.001105 |
