## Supplemental Table 3 for "Integrative Analysis the characterization of peroxiredoxins in pan-cancer"

Table S3 The associations between target drugs and PRDX expressions

| drug | PRDX1 | PRDX2 | PRDX3 | PRDX4 | PRDX5 | PRDX6 |
| --- | --- | --- | --- | --- | --- | --- |
| Vandetanib | 0.194647 | -0.0182 | 0.563382 | 0.339325 | 0.114883 | 0.438373 |
| Gemcitabine | 0.149144 | 0.070469 | 0.213025 | -0.05886 | -0.19928 | 0.051208 |
| AZD8055 | 0.147519 | 0.150818 | 0.084566 | 0.031925 | -0.04212 | 0 |
| Z-LLNle-CHO | 0.131679 | 0 | -0.06794 | 0.019206 | -0.03942 | 0.090389 |
| GDC-0941 | 0.131137 | 0.065384 | 0.317817 | 0.104295 | -0.04263 | 0.010344 |
| Doxorubicin | 0.128033 | 0.110062 | 0.179063 | -0.10524 | -0.17022 | 0.478478 |
| Irinotecan | 0.11119 | 0.094764 | 0.184622 | -0.12484 | -0.18223 | 0.041502 |
| 17-AAG | 0.107565 | 0 | 0.180897 | -0.04648 | 0.054854 | 0.091082 |
| lapatinib | 0.087885 | 0.036537 | 0.102143 | -0.14497 | 0.120357 | 0.086827 |
| Mitomycin-C | 0.08725 | 0.184876 | 0.104634 | -0.07691 | -0.12918 | 0.025398 |
| temsirolimus | 0.082543 | 0.145909 | 0.107624 | 0.264633 | -0.05428 | 0.363514 |
| Crizotinib | 0.081912 | 0.095706 | 0.319006 | 0.263822 | -0.07995 | 0.527947 |
| Pazopanib | 0.053164 | 0 | 0.200807 | 0.20384 | -0.0525 | 0.478478 |
| BIBR-1532 | 0.050339 | 0.182783 | 0.051799 | 0.012032 | -0.02248 | 0 |
| GNF-2 | 0.021132 | 0 | -0.04187 | -0.05813 | -0.09106 | 0.200791 |
| FTI-277 | 0.018747 | 0 | -0.0439 | -0.00509 | 0.082744 | 0.033056 |
| bortezomib | 0 | 0.167132 | 0 | 0 | 0 | 0 |
| TAE684 | -0.01805 | 0.089642 | -0.12224 | 0.154983 | 0.094301 | 0.077648 |
| OSI-906 | -0.03159 | 0.074328 | -0.06358 | 0.089355 | 0.076518 | -0.05009 |
| salermide | -0.04906 | 0.168997 | -0.03464 | 0.032716 | 0 | 0 |
| cediranib | -0.10654 | 0.04304 | 0.051591 | 0.118544 | -0.02912 | -0.08283 |
| BIX-01294 | -0.11153 | 0.18406 | -0.10008 | -0.09922 | -0.09112 | -0.17917 |
| staurosporine | -0.12849 | 0.005329 | 0.019075 | 0.174245 | -0.01122 | -0.06783 |
| Topotecan | -0.16783 | 0.160954 | 0.182319 | -0.08754 | -0.11819 | -0.05245 |
| ML312 | -0.18256 | 0.105382 | -0.07989 | 0.178524 | 0.01736 | -0.02497 |
