## Supplementary figures and images for "Integrative Analysis the characterization of peroxiredoxins in pan-cancer"

### Supplemental Figure1

#### Supplymentary Fig1

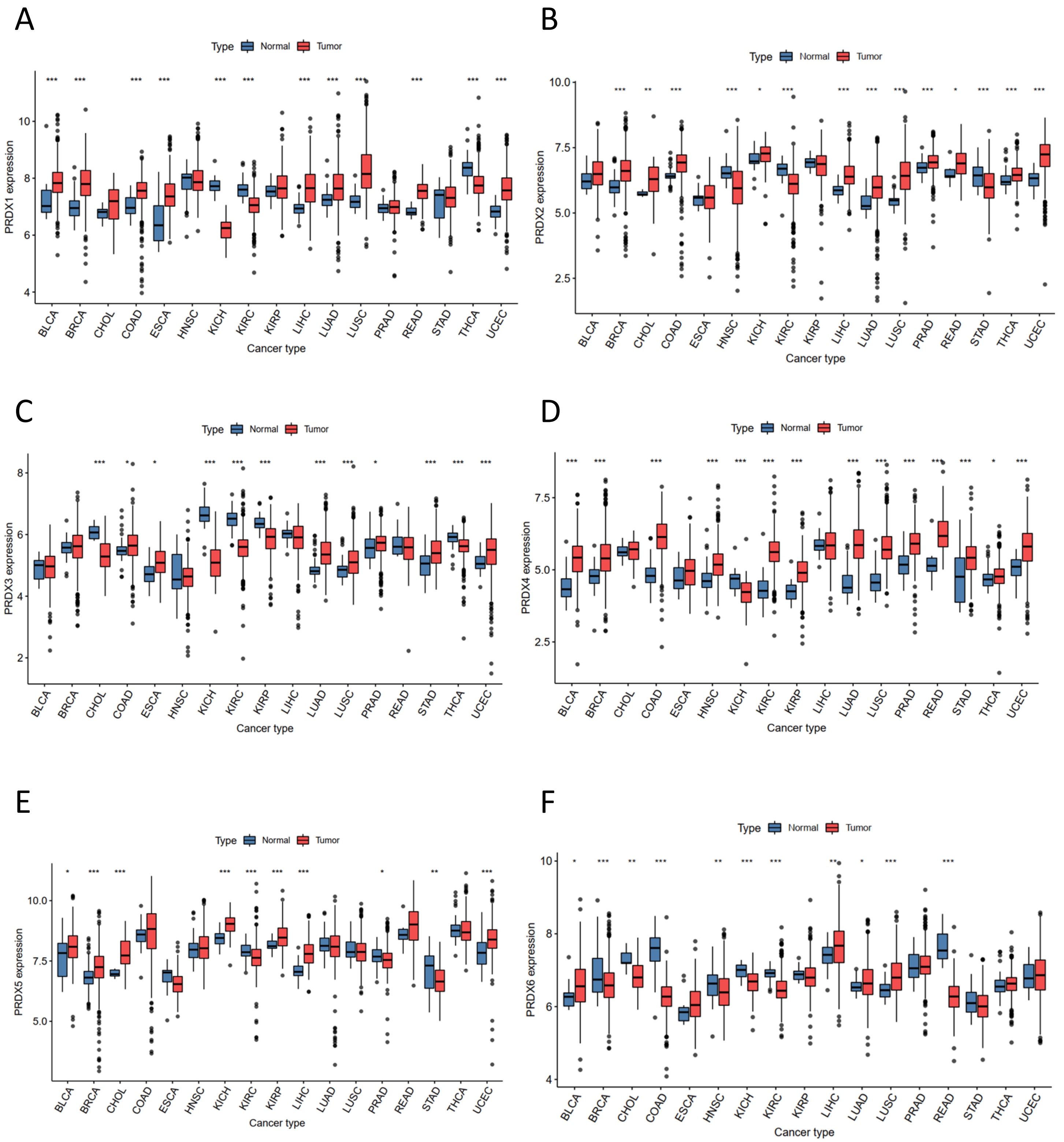

### Supplemental Figure2

Supplymentary Fig2

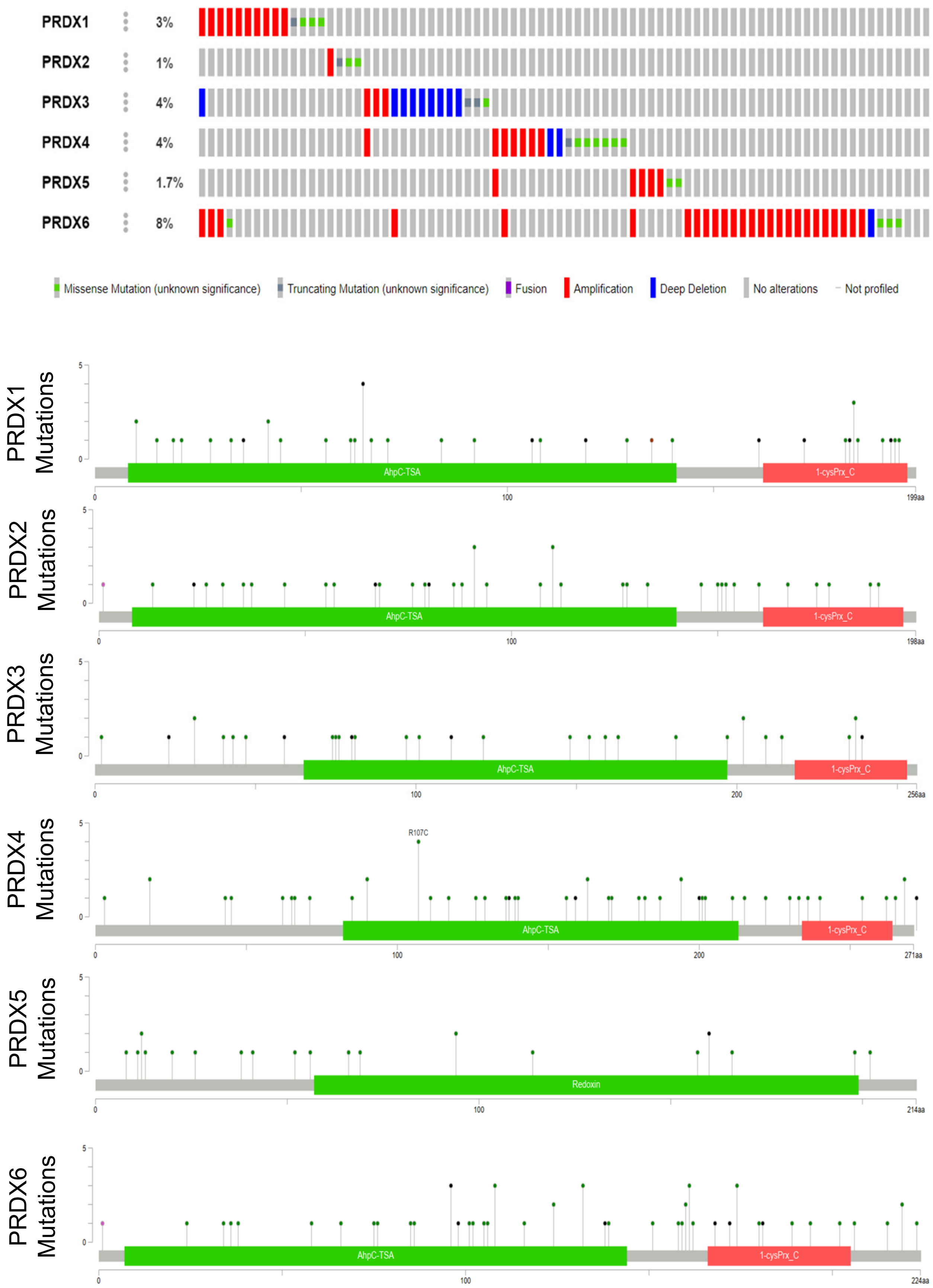
